# Reward Modality and Task Complexity Jointly Shape Learning Dynamics in Mice

**DOI:** 10.64898/2025.12.01.691606

**Authors:** Juliette Archimbaud, Charlotte Lehericy, Céline Auger, Christelle Rochefort, Laure Rondi-Reig

## Abstract

Learning efficacy relies on balancing reward-driven motivation with the difficulty of reaching a goal. Disentangling these factors is essential for understanding learning dynamics. Spatial navigation is a standard paradigm for probing these processes in rodents, but variation in mazes, cues, and protocols makes it difficult to determine how reward properties, rather than task design, shape learning and exploration,exploitation strategies. To date, no studies have directly compared reward type and learning difficulty within a single behavioral context.

Here, we used the Starmaze to examine how reward modality and task complexity jointly influence learning and the transition from exploration to exploitation. We created three versions of the same sequential egocentric task in which a fixed goal was rewarded with either a hidden escape platform (aquatic version), food pellets, or medial forebrain bundle stimulation. Task difficulty varied by increasing the number of required decision points from zero to three. Our results reveal distinct behavioral dynamics across reward modalities and provide a unified framework for studying reward-dependent learning in sequential navigation.

## Introduction

Spatial navigation refers to the ability to move around and find our way in various environments using either allocentric (world-centered) and/or egocentric (body-centered) representations and a variety of possible strategies, depending on the sensory information available to the navigator and the complexity of the task (Trullier et al., 1997; Arleo & Rondi-Reig, 2007; Vijayabaskaran et al., 2025). Spatial navigation relies on the combination of wider functions such as multi-sensory integration, motor control, learning and memory, as well as executive functions such as flexibility, to result in the production of a specific trajectory in a given context.

Although navigation has classically been investigated through simple tasks allowing to isolate discrete components of spatial cognition, such as the Radial Arm Maze for working memory (Olton & Samuelson, 1976), the Y-maze for cognitive flexibility (Havekes et al., 2006), or the openfield for spatial representation and exploration patterns (Walsh & Cummins, 1976, O’Keefe & Conway, 1978), these paradigms rarely engage these functions in isolation. Instead, animals must simultaneously integrate motivational, motor, and cognitive processes to perform goal-directed behaviors. This complexity makes it difficult to clearly dissociate the respective contributions of each process to task performance.

One promising framework to address this challenge is to interpret spatial navigation through the lens of the exploration–exploitation tradeoff, a behavioral dynamic well studied in decision-making and reinforcement learning (Cohen & Ranganath, 2007; Mehlhorn et al., 2015; Addicott et al., 2017). During early exploration, the animal samples the environment, identifies cues, and builds a representation of its surroundings. As information accumulates, behavior transitions to exploitation, where known paths and learned strategies are used to improve efficiency and goal-reaching performance. These two phases recruit distinct cognitive operations: exploration prioritizes novelty detection, uncertainty reduction, and behavioral flexibility, while exploitation relies on memory retrieval and efficient motor execution to get to the reward (Daw et al., 2006; Cinotti et al., 2019).

Furthermore, studying such complex behavior requires an equally sophisticated paradigm. The Starmaze, composed of a central pentagon with five radiating alleys, was originally developed to distinguish between allocentric and sequential egocentric navigation strategies in rodents (Rondi-Reig et al., 2006), and was later adapted for use in humans (Iglói et al., 2009; Maier et al., 2023). Its flexible structure allows for the selective opening or closing of specific alleys, enabling the precise manipulation of task complexity to assess its impact on learning and memory dynamics in a spatial context. Furthermore, the Starmaze can be implemented in both aquatic (Rondi-Reig et al., 2006) and dry environments (Cabral et al., 2014), providing a unique opportunity to compare how different motivational systems, such as escape from water or approach towards a food reward, affect navigational strategies and performance.

While both approaches have been widely used, they have however been associated with stress responses that may interfere with learning and memory performance (Harrison et al., 2009). An emerging alternative that emancipates from these confounding factors is the use of medial forebrain bundle (MFB) stimulation, a technique shown to elicit strong reward signals (Olds & Milner, 1954) and recently demonstrated to efficiently support the acquisition of a complex sequential foraging task (Zhang et al., 2023).

Here, we took advantage of the flexible design of the Starmaze to investigate how reward type and task complexity jointly shape learning performance and the exploration-exploitation dynamics.

In the current work, we created three versions of the same sequential egocentric task, each using a different reward type, located at the same position. These included: (i) locating a hidden platform in the aquatic version of the Starmaze (platform group), (ii) receiving a food pellet at the end of the path (food group), and (iii) receiving MFB stimulation at the goal location (stimulation group). We also manipulated task difficulty, from a simple path with no decision points to routes requiring one, two, or three correct turns to complete the correct trajectory (see Figure 1).

**Figure 1.**
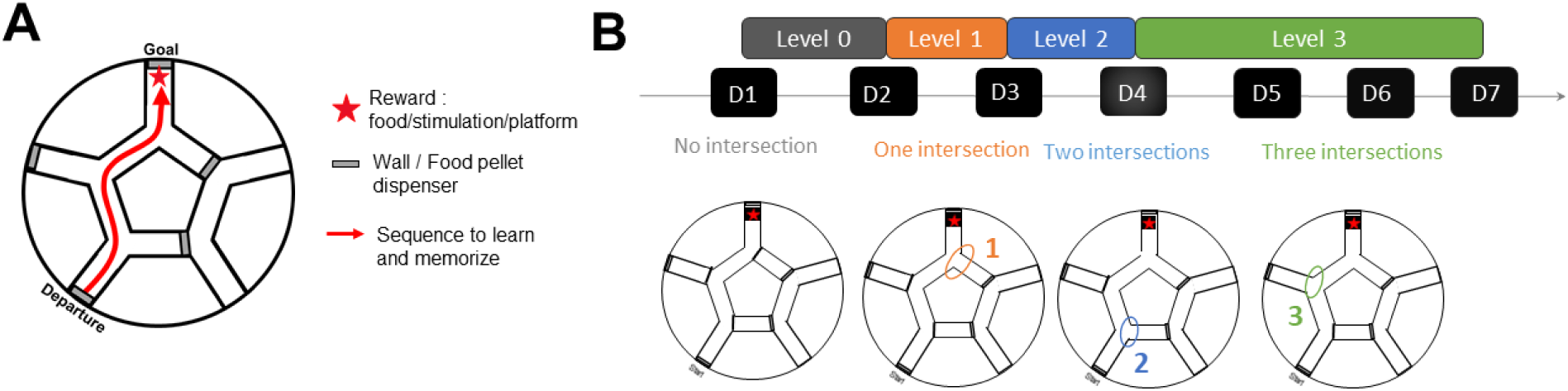
Diagram of the experimental setup. **(A)** Mice are trained in the Starmaze, a maze with five peripheral alleys. Training is conducted progressively, with intersections numbered 1, 2, and 3 being opened incrementally from day 1 to day 4. The sequence to be acquired is represented in red. Three groups of mice, each receiving a distinct type of reward (indicated by a red star), are involved in the experiment: a food reward, intracranial stimulation of the medial forebrain bundle (MFB), or access to a platform allowing the mice to escape from water. **(B)** The protocol lasts for 7 days (D1 to D7) and includes four levels of difficulty, denoted by numbers (0, 1, 2, and 3), which correspond to the number of peripheral alleys that are opened. Each day consists of 4 sessions (S), and each session comprises four trials.

This study provides a solid framework to assess how reward diversity and task complexity interact to shape the exploration–exploitation tradeoff and the dynamics of learning a sequential goal-directed navigation task.

## Results

To test for the efficiency of different types of reward, mice performances were evaluated in three different versions of the spatial navigation task, dry or aquatic, associated with a different type of reward delivery. Depending on the version of the task, mice had to reach the goal zone to be rewarded either with MFB stimulation (stimulation group, n=6), food (food group, n=14), or with the ability to escape from water on a dry platform (platform group, n=14). For the MFB group, the stimulation intensity was individually determined prior to training using a conditioned place preference test (see Supp. Fig. 1) and electrode placement was systematically verified post mortem (see supp Fig 2).

**Figure 2.**
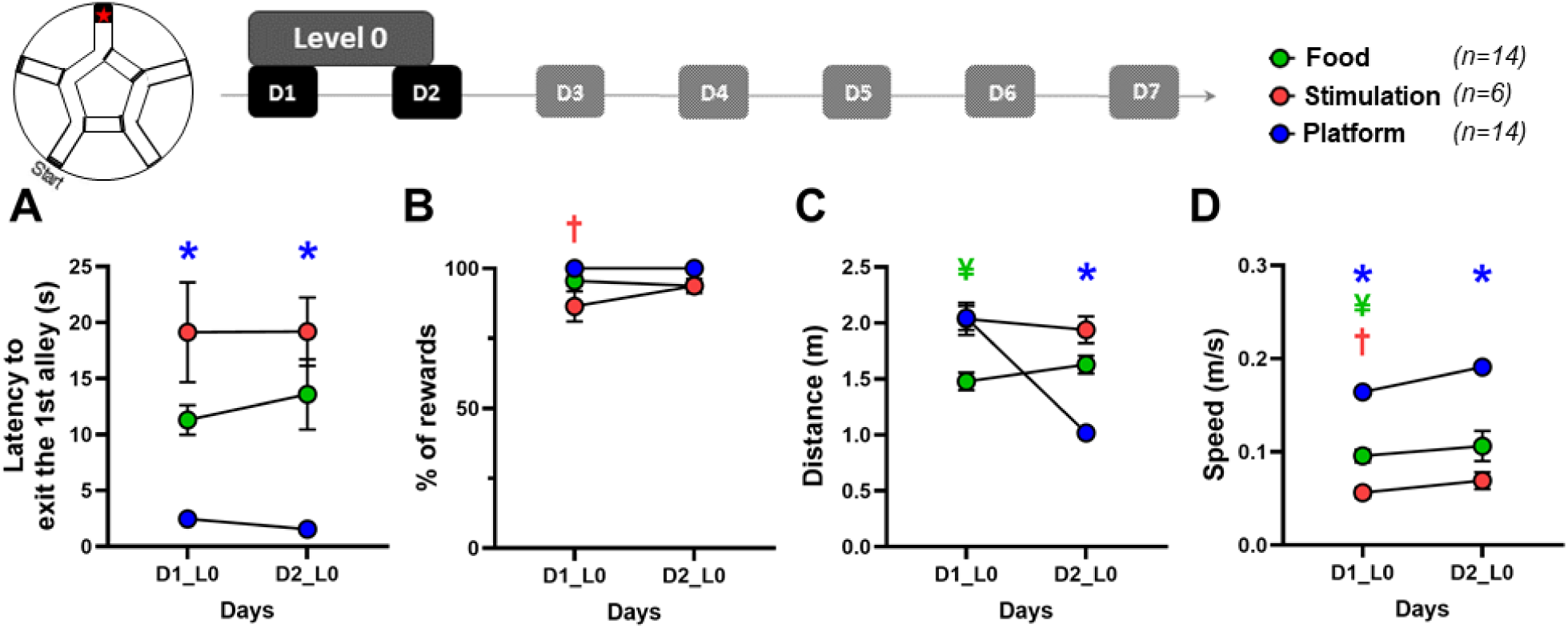
Goal-directed sequence with no decision point (level 0, “no intersection”). This figure illustrates the performances of the three groups (platform, stimulation, and food) compared in terms of A. latency to exit the first alley of the starmaze; B. percentage of trials in which mice get the reward; C. distance covered to reach the reward; D. the mean speed. *\* = the platform group is different from the two other groups. † = the stimulation group is different from the two other groups. ¥ = the food group is different from the two other groups (2-way repeated measures ANOVA with Tukey’s multiple comparisons tests)*.

### Motivational and performance differences between aquatic and land-based groups in the absence of decision points

We assessed several parameters reflecting willingness to engage in the task, performance, motivation, and path optimization across the different levels of difficulty of the starmaze sequence task. We first analyzed mice performances during the first two days of training, when the only available path in the maze led directly to the reward (see Figure 1). Under these conditions, with all intersections closed and no decision point, mice follow a unique route to get the reward. This phase allowed us to assess mice intrinsic motivation to get the reward without including any spatial processing or decision-making. Unexpectedly, MFB-rewarded mice showed slightly lower motivation compared to both the food- and platform-rewarded groups, as indicated by their longer latency to exit the start alley *(Fig. 2A, 2-way RM ANOVA, Day effect ns, Group effect p<0.0001, Day x Group ns, post-hoc Tukey’s multiple comparisons tests: Stimulation v Platform, p<0.0005, Stimulation v Food p=0.06*) and the lower percentage of rewards obtained on the first day (*Fig. 2B, 2-way RM ANOVA, Day effect ns, Group effect p<0.005, Day x Group ns, post-hoc Tukey’s multiple comparisons tests: Stimulation v Platform, p<0.0005, Stimulation v Food p<0.05*). Analysis of the distance traveled toward the reward and speed also revealed differences between the three groups of mice. The aquatic group showed a clear path optimization from day 1 to day 2, reaching the reward more rapidly. This improvement, reflected by both a reduction in distance (*Fig 2C, 2-way RM ANOVA, Day effect p<0.0005, Group effect p<0.0005, Day x Group p<0.0001, post-hoc Tukey’s multiple comparisons tests: D1 v D2, p<0.0005*, *Tukey’s multiple comparisons test*) and an increase in speed (*Fig 2D*, *2-way RM ANOVA, Day effect p<0.005, Group effect p<0.0001, Day x Group ns, post-hoc Tukey’s multiple comparisons test: D1 v D2, p<0.005*) was not observed in the other two groups (*D1 v D2, p>0.05* for both groups and both parameters).

In summary, at level 0 of training, when no memory of the route is required, mice in the aquatic condition exhibited better performances from the very first day of training with a higher speed, a lower latency to exit the departure alley, compared to both the MFB- and food-motivated groups.

### Increasing task difficulty reveals reward-specific patterns of navigation adaptation

We next investigated the dynamics of mice navigation under more challenging conditions, when mice were required to make one correct decision (turn left at the final junction to reach the reward, Figure 3 A-D) or two correct decisions (turn left at the first intersection and turn again left at the final junction, Figure 3 F-G). In such conditions, called Level 1 and Level 2, respectively, mice had to remember to turn left when a choice was required, both at the first and last turns.

**Figure 3.**
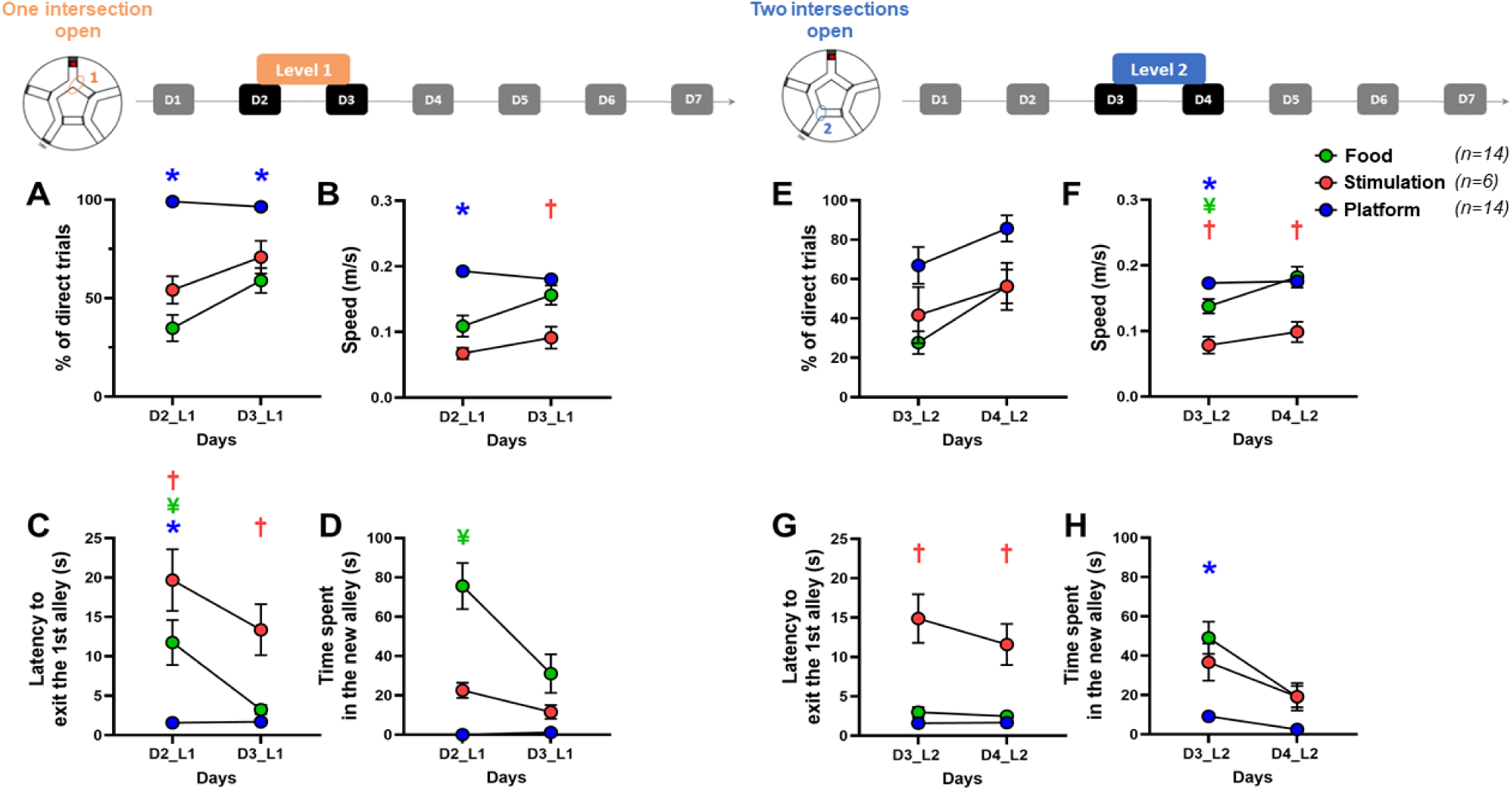
Goal-directed sequence with one (level 1) or two decision points (level 2). This figure compares the performances of the three groups (aquatic, MFB, and food) in terms of A. E. percentage of trials in which mice get directly to the reward; B. F. the mean speed; C. G. the latency to exit the departure alley; D. H. the time spent in the newly-opened alley. *\* = the platform group is different from the two other groups. † = the stimulation group is different from the two other groups. ¥ = the food group is different from the two other groups (2-way repeated measures ANOVA with Tukey’s multiple comparisons tests)*

This cognitive demand was supported differently by the three groups of mice. At Level 1, the aquatic-tested group used the optimal trajectory to the goal as soon as the first day of this new condition, outperforming the two land-based groups, as illustrated by the percentage of direct trials (*Fig. 3A, 2-way RM ANOVA, Day effect p<0.0005, Group effect p<0.0001, Day x Group p<0.0005, post-hoc Tukey’s multiple comparisons tests: D2_L1: Platform v Stimulation, p<0.0001, Platform v Food p<0.0001*) and speed (*Fig. 3B, 2-way RM ANOVA, Day effect p<0.0005, Group effect p<0.0001, Day x Group p<0.0001, post-hoc Tukey’s multiple comparisons tests: D2_L1: Platform v Stimulation, p<0.0001, Platform v Food p<0.0001*). Interestingly, although both MFB-stimulated and food-rewarded mice improved their trajectory over the two days of Level 1, they manifested different levels of motivation as illustrated by the latency to start the task (Fig 3C) and the exploratory behavior in the new alley of the Starmaze: while the MFB stimulated mice stayed longer in the departure alley before starting the task on both days (*Fig. 3C, 2-way RM ANOVA, Day effect p<0.0005, Group effect p<0.0001, Day x Group p<0.005, post-hoc Tukey’s multiple comparisons tests: D2_L1: Stimulation v Food, p<0.05, Stimulation v Platform p<0.0001, D3_L1: Stimulation v Food, p<0.005, Stimulation v Platform p<0.005*), the food rewarded group spent more time in the new alley on the first day (*Fig. 3D, 2-way RM ANOVA, Day effect p<0.005, Group effect p<0.0001, Day x Group p<0.005, post-hoc Tukey’s multiple comparisons tests: D2_L1: Food v Stimulation, p<0.005, Food v Platform p<0.0001*) before reaching stimulated mice level on Day 2, remaining still higher than the aquatic group (*Fig. 3D, post-hoc Tukey’s multiple comparisons tests: D3_L1: Food v Platform p<0.05*).

The addition of another decision point at Level 2 induced contrasting results in the three conditions tested regarding mice capacity to adapt their trajectory. While the aquatic group performances were mildly affected by the presence of a second intersection, they remained at a superior level in terms of path optimization as shown by the increased percentage of direct trajectory (*Fig. 3E, 2-way RM ANOVA, Day effect p<0.0005, Group effect p<0.005, Day x Group ns, post-hoc Tukey’s multiple comparisons tests: Platform: D3_L2 v D4_L2, p<0.05*), highest speed (*Fig. 3F, 2-way RM ANOVA, Day effect p<0.0005, Group effect p<0.0001, Day x Group p<0.005, post-hoc Tukey’s multiple comparisons tests:D3_L2: Platform v Stimulation, p<0.0001, Platform v Food p<0.05*) and the lowest time spent in the new alley (*Fig. 3H, 2-way RM ANOVA, Day effect p<0.05, Group effect p<0.0001, Day x Group p<0.05, post-hoc Tukey’s multiple comparisons tests:D3_L2: Platform v Stimulation, p<0.05, Platform v Food p<0.0001*). Remarkably, the food tested mice showed a strong improvement in all the trajectory-associated parameters over the two days of Level 2 as reflected by the increased percentage of direct trials (*Fig. 3E, Tukey’s multiple comparisons tests: Food: D3_L2 v D4_L2, p<0.0005*), speed (*Fig. 3F, Tukey’s multiple comparisons tests: Food: D3_L2 v D4_L2, p<0.0001*) as well as the reduced time spent in the new alley (*Fig. 3H, Tukey’s multiple comparisons tests: Food: D3_L2 v D4_L2, p<0.0001*). On the other hand, MFB-stimulated mice didn’t significantly optimize their trajectory as reflected by the percentage of direct trials, even though they reduced their exploration of new peripheral alleys (*Fig. 3H, Tukey’s multiple comparisons tests: Stimulation: D3_L2 v D4_L2, p<0.05*). While their latency to exit the first alley decreased between the days of level 2 (*Fig. 3G, Tukey’s multiple comparisons tests: D3_L2 v D4_L2, p<0.005*), this motivation to initiate the task still remained lower than the two other groups (*Fig. 3G, Tukey’s multiple comparisons tests: D3_L2 and D4_L2: Stimulation v Platform, p<0.0001, Stimulation v Food p<0.0001*).

### High cognitive demand equalizes performance through reward-dependent strategies

Finally, we investigated mice performance dynamics at Level 3, when the cognitive demand increases to a full sequence of three consecutive correct turns to remember (left-right-left). Given the complexity of the task, we evaluated the performance across four days of training from Day 4 to Day 7 (Figure 4). In this task level, the three groups showed equal ability to optimize their path to get the reward in terms of both performance quality and dynamics, as shown by the percentage of direct trials *(Fig. 4A, 2-way RM ANOVA, Day effect p<0.0001, Group effect ns, Day x Group ns*). To reach such performance levels, the three groups appeared to rely on different strategies, as reflected by changes in other behavioral parameters. Unlike the aquatic group, land-based tested mice (stimulation + food) showed an increase in speed over days (*Fig. 4B, 2-way RM ANOVA, Day effect p<0.0001, Group effect p<0.0001, Day x Group p<0.0001, post-hoc Tukey’s multiple comparisons tests: Food: D4_L3 v D7_L3 p<0.0001, Stimulation: D4_L3 v D7_L3 p<0.05*), with the most pronounced improvement observed in the food-rewarded group, which reached a significantly higher speed than the other groups by the end of training (*Fig. 4B, Tukey’s multiple comparisons tests: D7_L3: Food v Stimulation, p<0.005, Food v Platform p<0.05*). Interestingly, mice in the MFB group exhibited lower goal-directed motivation and a greater exploratory behavior upon exposure to novelty, as respectively indicated by the tendency to spend more time in the departure alley (*Fig. 4C, 2-way RM ANOVA, Day effect p<0.0005, Group effect p<0.0001, Day x Group p<0.0001*) and more time in the new alley at the onset of Level 3 (*Fig.4D, 2-way RM ANOVA, Day effect p<0.0001, Group effect p<0.005, Day x Group p<0.005*).

**Figure 4.**
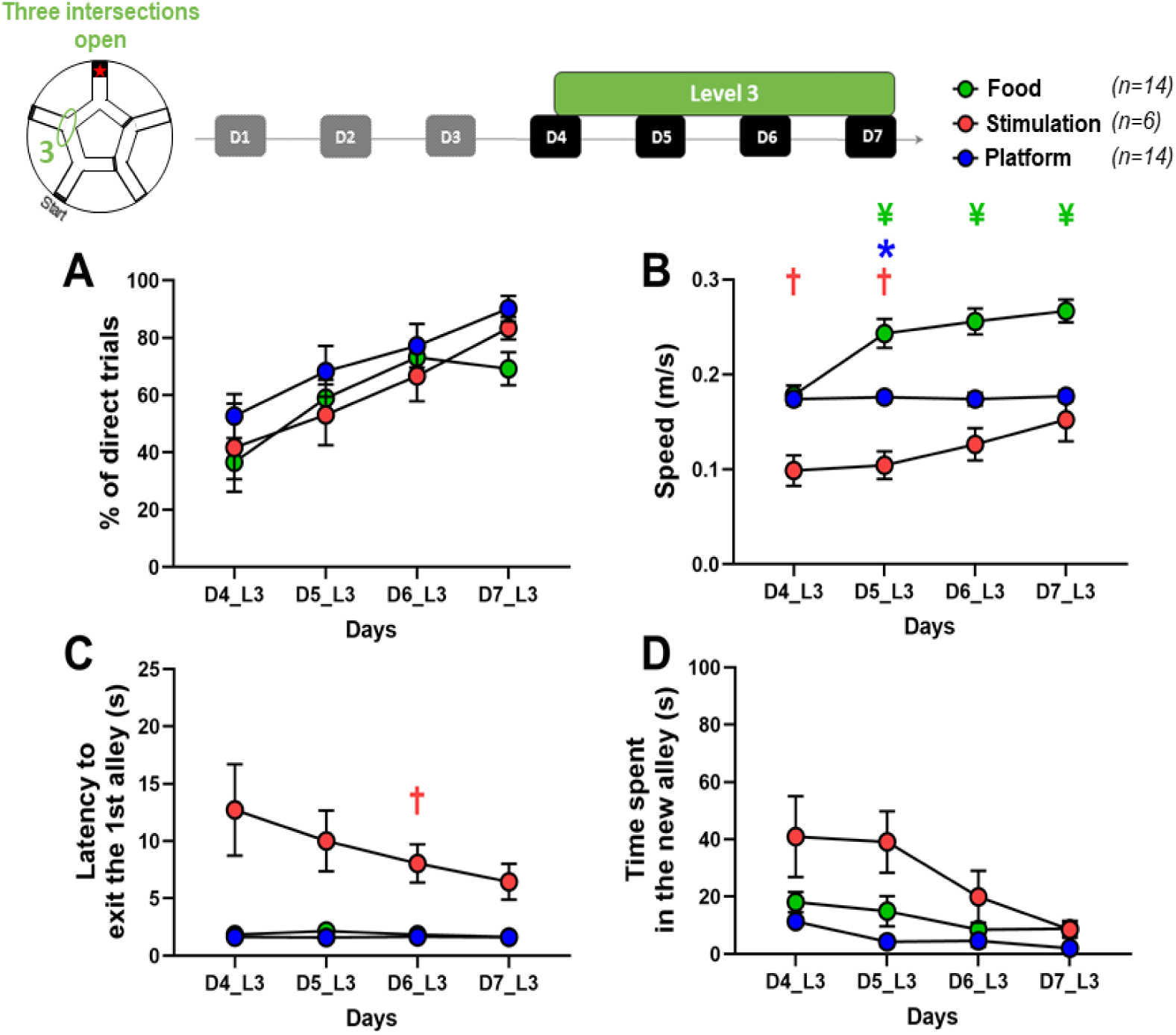
Level 3 (A 3-turn sequence to remember). This figure illustrates the percentage of direct trials (A), speed (B), latency to exit the first alley (C), and time spent in the newly-opened arm (D) for level (L) 3, where all peripheral alleys are open. The metrics are displayed across days (D4 to D7) for three groups of mice receiving different types of rewards: food (green), MFB stimulation (red), and a platform for escaping water (blue).**\* = the platform group is different from the two other groups. † = the stimulation group is different from the two other groups. ¥ = the food group is different from the two other groups** *(2-way repeated measures ANOVA with Tukey’s multiple comparisons tests)*

Overall, as task complexity increased, the sharp distinction between aquatic and land-tested groups gave way to more subtle differences, with performance levels converging across all three groups. To further disentangle the strategies supporting this apparent convergence in performance, we next examined how trajectory optimization evolved on a finer timescale by analyzing distance traveled across sessions from Day 4 to Day 7.

### Trajectory optimization relies on reward-modulated memory dynamics

To further explore learning of a complex sequence in the three conditions tested, we used distance-based scores reflecting different mnesic components. First, we looked at memory consolidation overnight by comparing the distance animals traveled in the first session of a day with the last session of the previous day (*Fig. 5B*). Only the aquatic group showed efficient memory consolidation during Level 3 with a distance travelled during S1 of each day that was lower that during S4 of the previous day (*Fig. 5B, Platform: p<0.05 for D4_L3-D5_L3 and D6_L3-D7_L3, One sample Wilcoxon test)*. In contrast, the land-based groups, especially those rewarded with food, consistently began each day by covering more distance than the previous one, although this difference gradually diminished over time (*Fig. 5B, Food: p<0.05 for D4_L3-D5_L3 and D5_L3-D6_L3, One-sample Wilcoxon test)*. Secondly, we measured memory within a day by comparing the distance traveled between the last and first session of each day (*Fig. 5C*). There were no clear changes in this parameter over days in the aquatic group. However, the food group showed a strong ability to correct their trajectory as reflected by the reduced distance during the last session compared to the first (*Fig. 5C, Food: p<0.05 for D5_L3 and D6_L3, One-sample Wilcoxon test)*. The MFB-stimulated group, on the other hand, showed improvement as seen by the difference in traveled distance between the last and first session of Day 5, with this gap decreasing over time and aligning with the aquatic group by Day 6 (*Fig. 5C*).

**Figure 5.**
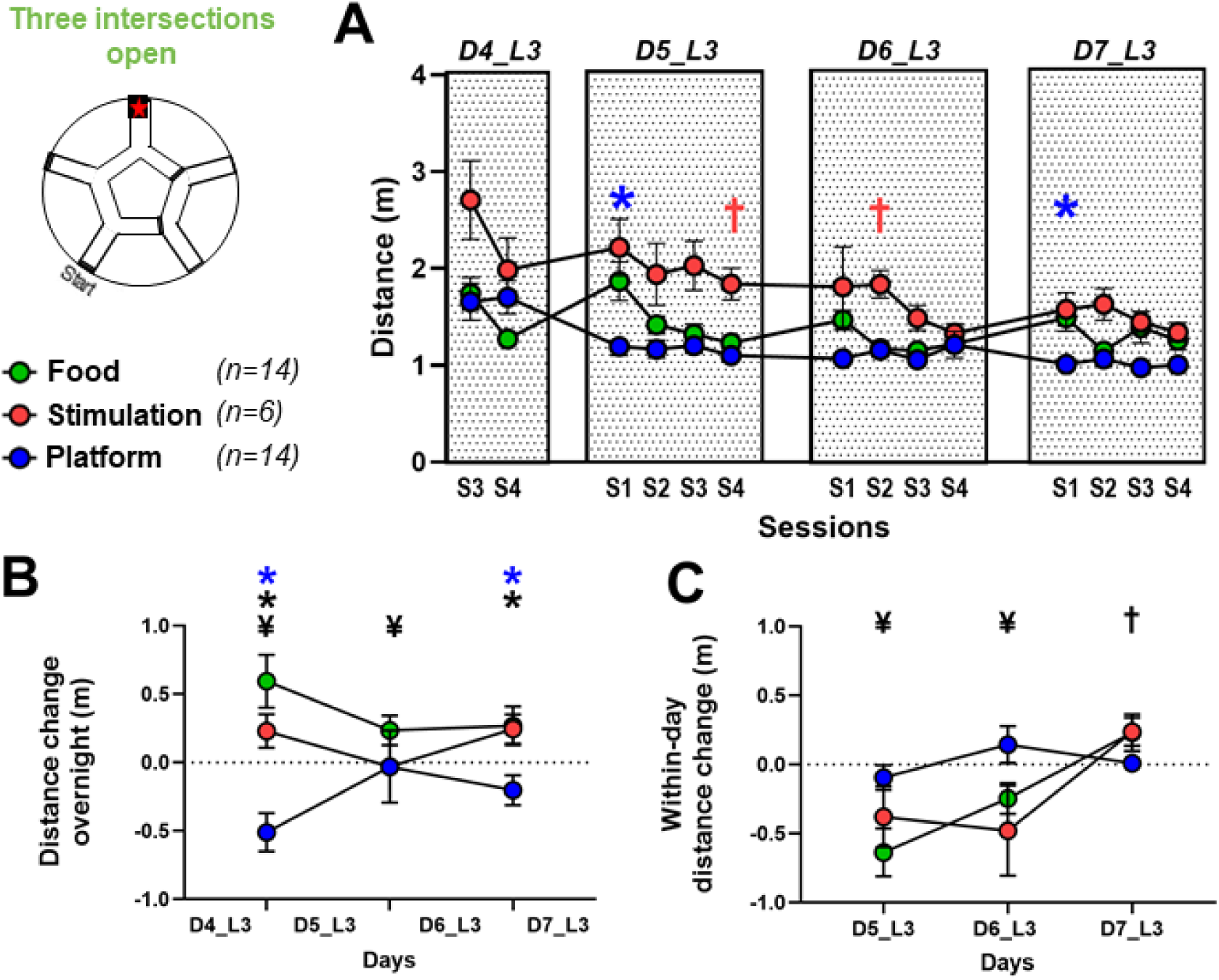
Dynamics of path optimization at Level 3. This figure illustrates the (A) distance, (B) distance difference between the last session of a day and the first session of the following day (S1_n_-S4_n-1_), and (C) difference between distance travelled at the first and last session of each day (S4-S1) of the Level 3, where all peripheral alleys are open. The metrics are displayed across Sessions (A) or days (B, C) (D4 to D7) for three groups of mice receiving different types of rewards: food (green), MFB stimulation (red), and a platform for escaping water (blue). Negative values for (B) and (C) indicate a reduction in the distance between two consecutive sessions.***\* = the aquatic group is different from the two other groups. † = the MFB group is different from the two other groups. ¥ = the food group is different from the two other groups*** *(2-way repeated measures ANOVA with Tukey’s multiple comparisons tests). Symbols in black indicate a score significantly different than 0 (One-sample Wilcoxon tests)*

In summary, although overall performance levels were similar between the three groups, trajectory analyses on a finer scale revealed distinct dynamics during learning of a complex three-turn sequence: aquatic-tested mice rapidly optimized their path and maintained stable performance, food-rewarded mice showed fluctuating trajectories and within-day correction, and MFB-stimulated mice exhibited a gradual and delayed improvement.

## Discussion

In this study, we compared the effects of three distinct rewards (escaping water, getting food, or receiving a rewarding electrical stimulation) within a standardized learning protocol in the Starmaze while varying the task complexity. This approach enabled us to investigate how reward sources (internal vs external), valence (negative vs positive reinforcers), stressors (external as water immersion vs internal as hunger), and cognitive load shape not only task performance but also the dynamics of learning as well as the balance between exploration and exploitation.

Earlier studies comparing aquatic and dry mazes have produced conflicting results: while some reports suggest that animals learn better in aquatic environments (Hodges, 1996), others report equal or even better performances in land-based setups (Patil et al., 2009; Llano Lopez et al., 2010). However, these controversial findings often stem from tasks performed across different mazes (i.e., radial arm maze versus Morris Water Maze), making direct comparisons difficult. To our knowledge, our design allows for the first time a direct comparison of aquatic and several dry reward conditions in an identical navigation task, providing new insights into how reward properties influence both performance and learning dynamics.

In line with findings of Ormerod et al. (2002), who reported that learning was faster in water than on land in a delayed match-to-position task, the aquatic group shifted from exploration to exploitation earlier than the two land-based groups. The study from Hodges (1996) similarly noted that rats performed more consistently in the Morris Water Maze compared to dry-land setups. This pattern suggests that aversive reinforcement, rather than positive one, has a greater impact on learning efficacy in a spatial context. Prior studies have shown that rodents receiving MFB stimulation as reward outperform those subjected to food or water deprivation in operant or perceptual tasks (McMurray et al., 2017; Verdier et al., 2022). Surprisingly, our data indicate that, in a spatial navigation task, MFB-stimulated mice consistently displayed longer latencies before engaging with the maze. This discrepancy cannot be attributed to stimulation inefficacy, as the parameters were individually validated via a conditioned place preference protocol prior to the Starmaze task (supplementary Fig. 1). Therefore, it is possible that MFB stimulation interacts differently with complex goal-directed spatial behaviors than with simpler operant or perceptual tasks, and that the absence of deprivation may delay initial engagement and higher exploration of new alleys in a foraging-like environment. Notably, MFB stimulation does not engage physiological deprivation systems (e.g., hunger or thirst), which are known to drive immediate exploratory behavior in foraging contexts. Sasaki et al. (1984) hypothesized that electrical stimulation of the MFB can mimic the rewarding effect of food deprivation. However, they demonstrated that although MFB stimulation and natural rewards both inhibit the same lateral hypothalamic neurons, their behavioral expressions are not always equivalent, particularly in tasks requiring goal-directed navigation.

### Task complexity modulates the relative contribution of behavioral drivers

Increasing task complexity by introducing two decision points to perform the path (level 2) did not reveal differences in cognitive performance but rather uncovered differences in behavioral strategy across groups. This effect was particularly pronounced in the MFB-stimulated mice compared to the two other groups, as it presents longer onset latencies combined with reduced speed. Two non-mutually exclusive explanations may account for this difference with the two other groups: (1) the presence of stressors, whether external (water immersion) or internal (hunger), may enhance the urgency to obtain the reward in aquatic and food-rewarded mice; and (2) the physical and multi-sensory nature of the escape platform or food reward, in contrast to the virtual nature of MFB stimulation directly targeting brain reward circuit without any external sensory input, may facilitate a stronger and faster association between spatial location and reward. In line with this, Burnett et al. (2016) showed that physiological hunger can suppress competing motivational drives, which supports the idea that food-based reward may provide a more dominant framework for driving efficient spatial behavior than MFB stimulation. These findings suggest that as task complexity increases, the influence of stress and tangible, sensory-driven rewards becomes more critical in shaping spatial behavior than the environment and reward valence.

### Reward type influences strategy without affecting performance under high cognitive demand

At the highest level of task complexity (3 consecutive intersections to remember, level 3), all three groups—regardless of reward type—ultimately achieved similar learning outcomes, as reflected by progressively more direct trajectories over time. However, the timing of their transition from exploration to exploitation varied, indicating that distinct behavioral strategies were still present. While precisely defining the moment of this shift remained challenging, the combination of performance measures and motivation-related parameters enabled us to approximate when each group switched its behavioral mode.

For the mice navigating the aquatic starmaze, a rapid transition from exploration to exploitation was evidenced by the sharp drop of the exploration-related measures, such as time spent in novel alleys and distance travelled between the first and second day. This early behavioral shift, followed by steady path optimization in later sessions, suggests that these animals quickly acquired the goal location and then focused on refining their trajectory, indicating ongoing learning and spatial memory engagement rather than automatized responses. Importantly, this group was the only one to show significant day-to-day improvements in the traveled distance, indicating that overnight memory consolidation may have supported their learning process. These findings suggest that negative reinforcement through acute exposure to an external stressor, such as water immersion, can induce quick and efficient spatial learning and promote mechanisms of memory consolidation.

In contrast, food-rewarded animals did not exhibit such a clear behavioral shift but instead showed a more complex dynamic at this stage of the task. While exploratory behavior, reflected by time spent in new alleys, gradually declined between Days 4 and 6, a marked increase in speed between Days 4 and 5 suggested a rise in exploitation behavior. However, when analyzing path distance, these animals consistently began each day with poorer performance than the previous day, indicating a lack of strong memory consolidation overnight. Instead, performance typically improved between sessions within a single day, pointing to a within-session adjustment rather than sustained memory across days. Surprisingly, despite its previously described strong rewarding effects during spatial navigation (Kobayashi et al., 1997; Zhang et al., 2023), MFB stimulation led here to the most gradual learning profile, with a progressive evolution of behavior across Level 3. Improvements in speed and time spent in novel alleys emerged late (days 5–6), without any clear indication of a consolidation process across days or within sessions. A potential transition appeared on day 6, marked by a reduced distance before reaching a plateau. These results point to a slow, incremental learning process, in which the absence of aversive drive and the use of a virtual reward favor delayed engagement and a late shift from exploration to exploitation under high cognitive demand.

### Reward Diversity as a Tool to Dissect Neural Circuits Underlying Memory and Decision-Making Processes

Overall, our results demonstrate that sequence-based navigation can be efficiently learned using either positive or negative reinforcements, physical or virtual reward, even under increasing cognitive demand. However, the distinct behavioral strategies observed across conditions suggest the engagement of different, though potentially overlapping, neural circuits underlying learning and memory of a complex spatial sequence. The aquatic paradigm proved particularly useful for studying memory consolidation, as the strong performance and rapid trajectory refinement implied overnight integration of spatial information. Yet, due to the urgency imposed by the aversive context, aquatic trials offer limited resolution on decision-making at individual choice points.

In contrast, land-based protocols, particularly the food-rewarded condition, allowed more detailed analysis of moment-to-moment decision-making and working memory, as the lower cost of errors encouraged exploratory behavior. While these animals did not show strong consolidation across days, they exhibited within-day improvements, making this protocol especially suited to probing the dynamics of short-term memory and neural activity during decision-making in a spatial context. The MFB stimulation, despite slower optimization, offers a non-deprivation-based alternative paradigm that eliminates hunger-induced stress as a confounding factor. The gradual improvement of this group may reflect greater difficulty in forming spatial associations without physical cues, rather than a lack of motivation, potentially involving distinct circuitry related to planning or internal valuation processes.

The behavioral differences observed between the three reward conditions (aquatic, food, and MFB stimulation) suggest that distinct neural networks are differentially engaged depending on the motivational context, with potential consequences for the balance between exploration and exploitation. In the aquatic maze, escaping water likely induces a stress response leading to a rapid shift toward stable exploitation, involving not only a hippocampus-centered spatial memory network but also the dorsal striatum, which supports well-learned sequential strategies (Packard & McGaugh, 1996). In contrast, food reward was associated with greater initial exploration as illustrated by level 2 performances. This could reflect a preferential recruitment of hippocampal–prefrontal networks that promote cognitive flexibility and decision making. Finally, MFB stimulation, which directly activates mesolimbic and mesocortical dopaminergic pathways (Wise, 2004; Ilango et al., 2014), may involve ventral striatal and prefrontal networks, sustaining prolonged exploration and delaying the exploitation mode relative to the two other reward conditions. These findings raise the hypothesis that formally identical task demands can be supported by partly non-overlapping neural substrates depending on reward properties. Further investigation is required to test this hypothesis, and our Starmaze protocol offers a powerful tool to address this question by combining the current behavioral framework with neural recording or imaging approaches to map the recruitment and interaction of specific brain regions during sequence learning under different motivational contexts.

By leveraging rewards that vary in valence, tangibility, and associated stressors, our study reveals how these behavioral drivers shape learning strategies. These findings lay the groundwork for future investigations into the neural mechanisms of motivation, decision-making, and memory processes, including memory consolidation, across different cognitive and emotional contexts.

**Supp. Fig 1:**
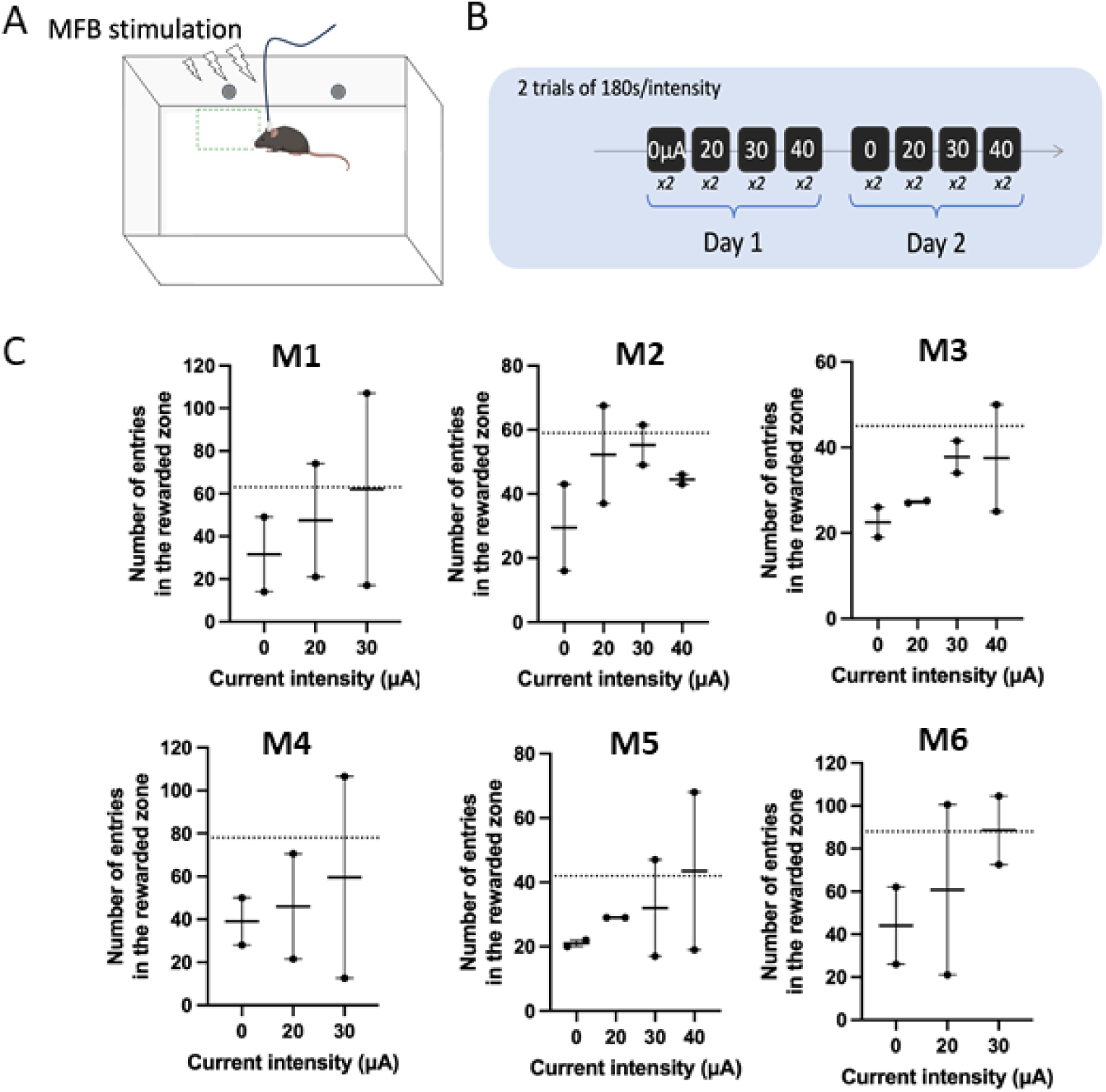
Functional validation of the MFB electrode implantation in the place preference task. **(A)** A train of MFB stimulation pulses was delivered upon entrance in the rewarded zone (green). **(B)** Place preference was tested two consecutive days at increasing stimulation current intensities from 0 to 40 µA (2 consecutive sessions of 180 s). **(C)** Number of entries into the rewarded zone as a function of current intensity for the 6 individual mice included in the MFB group (M1 to M6). Electrode implantation was validated if the number of entries in the rewarded zone was at least once doubled compared to the mean value at 0 µA (dotted line). MFB stimulation was functional for all 6 mice.

**Supp. Fig. 2:**
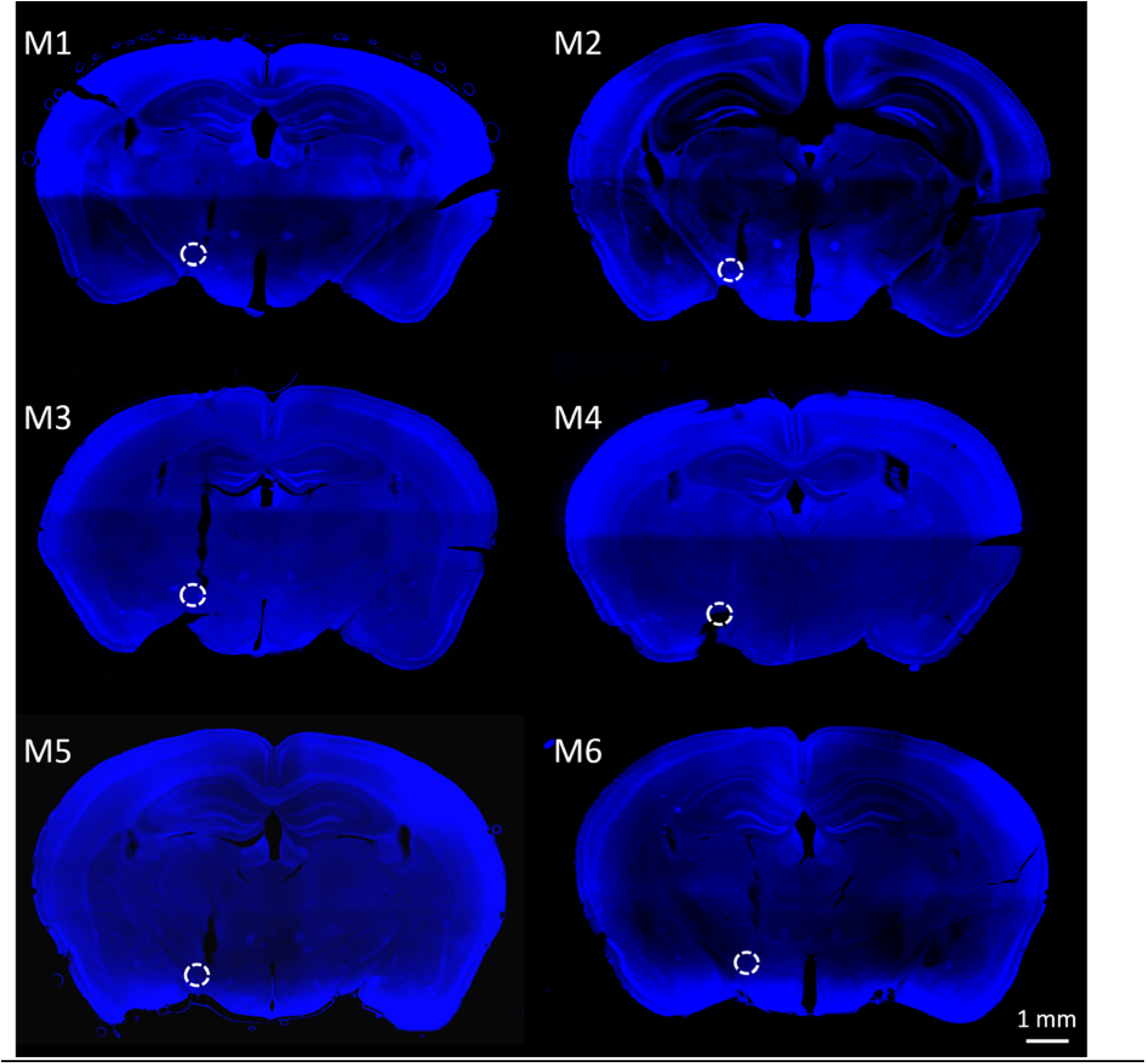
Histological validation of MFB electrode implantation. Representative coronal sections from the six mice of the MFB group (M1 to M6). White circles indicate electrode tip location. Sections were counterstained with DAPI.

**Supplementary Table 1:** Statistical analyses for all levels performances across the three reward conditions (MFB stimulation, food, and platform escape). The table reports 2-way repeated-measures ANOVAs (Group × Day) for four behavioral measures followed by Tukey’s post-hoc comparisons between groups for each testing day. For each contrast, q-values and adjusted P-values (Padj) are provided.

| Level 0 |  |  |  |  |
| --- | --- | --- | --- | --- |
| Latency to exit the first alley<br>2-way RM ANOVA | F(DFn, DFd) | P-value | Tukey's post hoc D1_L0 | Tukey's post hoc D2_L0 |
| Group effect | F (2, 31) = 24.36 | P<0.0001 | Platform v Stimulation<br>q=6.952 ; Padj<0.0001 | Platform v Stimulation<br>q=7.358 ; Padj<0.0001 |
| Day effect | F (1, 31) = 0.09055 | P=0.7655 | Platform v Food<br>q= 4.756 ; Padj=0.0037 | Platform v Food<br>q= 6.481 ; Padj<0.0001 |
| Interaction | F (2, 31) = 0.5069 | P=0.6073 | Stimulation v Food<br>q= 3.268 ; Padj=0.0617 | Stimulation v Food<br>q= 2.338 ; Padj=0.2313 |
| Percentage of rewards<br>2-way RM ANOVA | F(DFn, DFd) | P-value | Tukey's post hoc D1_L0 | Tukey's post hoc D2_L0 |
| Group effect | F (2, 31) = 7.601 | P=0.0021 | Platform v Stimulation<br>q=5.760 ; Padj=0.0004 | Platform v Stimulation<br>q=2.658 ; Padj=0.1531 |
| Day effect | F (1, 31) = 1.504 | P=0.2293 | Platform v Food<br>q= 2.451 ; Padj=0.2010 | Platform v Food<br>q= 3.432 ; Padj=0.0471 |
| Interaction | F (2, 31) = 2.712 | P=0.0821 | Stimulation v Food<br>q= 3.861 ; Padj=0.0221 | Stimulation v Food<br>q= 0.000 ; Padj>0.9999 |
| Distance<br>2-way RM ANOVA | F(DFn, DFd) | P-value | Tukey's post hoc D1_L0 | Tukey's post hoc D2_L0 |
| Group effect | F (2, 31) = 10.86 | P=0.0003 | Platform v Stimulation<br>q=0.07982 ; Padj=0.9982 | Platform v Stimulation<br>q=8.989 ; Padj<0.0001 |
| Day effect | F (1, 31) = 17.84 | P=0.0002 | Platform v Food<br>q=7.136 ; Padj<0.0001 | Platform v Food<br>q=7.680 ; Padj<0.0001 |
| Interaction | F (2, 31) = 29.91 | P<0.0001 | Stimulation v Food<br>q=5.447 ; Padj=0.0008 | Stimulation v Food<br>q=3.040 ; Padj=0.0883 |
| Speed<br>2-way RM ANOVA | F(DFn, DFd) | P-value | Tukey's post hoc D1_L0 | Tukey's post hoc D2_L0 |
| Group effect | F (2, 31) = 41.82 | P<0.0001 | Platform v Stimulation<br>q=9.655 ; Padj<0.0001 | Platform v Stimulation<br>q=10.87 ; Padj<0.0001 |
| Day effect | F (1, 31) = 9.452 | P=0.0044 | Platform v Food<br>q=7.908 ; Padj<0.0001 | Platform v Food<br>q=9.755 ; Padj<0.0001 |
| Interaction | F (2, 31) = 1.134 | P=0.3348 | Stimulation v Food<br>q=3.529 ; Padj=0.0399 | Stimulation v Food<br>q=3.315 ; Padj=0.0571 |

| Level 1 |  |  |  |  |
| --- | --- | --- | --- | --- |
| Percentage of direct trials<br>2-way RM ANOVA | F(DFn, DFd) | P-value | Tukey's post hoc D2_L1 | Tukey's post hoc D3_L1 |
| Group effect | F (2, 31) = 36.79 | P<0.0001 | Platform v Stimulation | Platform v Stimulation |
|  |  |  | q=7.319 ; Padj<0.0001 | q=4.168 ; Padj=0.0124 |
| Day effect | F (1, 31) = 18.38 | P=0.0002 | Platform v Food | Platform v Food |
|  |  |  | q= 13.52 ; Padj <0.0001 | q= 7.884 ; Padj<0.0001 |
| Interaction | F (2, 31) = 10.19 | P=0.0004 | Stimulation v Food | Stimulation v Food |
|  |  |  | q= 3.151 ; Padj=0.0744 | q= 1.939 ; Padj=0.3623 |
| Speed<br>2-way RM ANOVA | F(DFn, DFd) | P-value | Tukey's post hoc D2_L1 | Tukey's post hoc D3_L1 |
| Group effect | F (2, 31) = 17.45 | P<0.0001 | Platform v Stimulation | Platform v Stimulation |
|  |  |  | q=8.959 ; Padj<0.0001 | q=6.371 ; Padj<0.0001 |
| Day effect | F (1, 31) = 18.82 | P=0.0001 | Platform v Food | Platform v Food |
|  |  |  | q= 7.719 ; Padj<0.0001 | q= 2.227 ; Padj=0.2642 |
| Interaction | F (2, 31) = 21.00 | P<0.0001 | Stimulation v Food | Stimulation v Food |
|  |  |  | q= 2.980 ; Padj=0.0966 | q= 4.646 ; Padj=0.0047 |
| Latency to exit the first alley<br>2-way RM ANOVA | F(DFn, DFd) | P-value | Tukey's post hoc D2_L1 | Tukey's post hoc D3_L1 |
| Group effect | F (2, 31) = 16.71 | P<0.0001 | Platform v Stimulation | Platform v Stimulation |
|  |  |  | q=8.586 ; Padj<0.0001 | q=5.526 ; Padj=0.0007 |
| Day effect | F (1, 31) = 19.25 | P=0.0001 | Platform v Food | Platform v Food |
|  |  |  | q=6.234 ; Padj=0.0001 | q=0.9350 ; Padj=0.7868 |
| Interaction | F (2, 31) = 7.557 | P=0.0021 | Stimulation v Food | Stimulation v Food |
|  |  |  | q=3.757 ; Padj=0.0267 | q=4.802 ; Padj=0.0034 |
| Time spent in the new alley<br>2-way RM ANOVA | F(DFn, DFd) | P-value | Tukey's post hoc D2_L1 | Tukey's post hoc D3_L1 |
| Group effect | F (2, 31) = 22.20 | P<0.0001 | Platform v Stimulation | Platform v Stimulation |
|  |  |  | q=2.465 ; Padj=0.1975 | q=1.137 ; Padj=0.7019 |
| Day effect | F (1, 31) = 9.494 | P=0.0043 | Platform v Food | Platform v Food |
|  |  |  | q=10.67 ; Padj<0.0001 | q=4.228 ; Padj=0.0110 |
| Interaction | F (2, 31) = 7.509 | P=0.0022 | Stimulation v Food | Stimulation v Food |
|  |  |  | q=5.803 ; Padj=0.0004 | q=2.138 ; Padj=0.2925 |

| Level 2 |  |  |  |  |  |
| --- | --- | --- | --- | --- | --- |
| Percentage of direct trials<br>2-way RM ANOVA | F(DFn, DFd) | P-value | Tukey's post hoc<br>D3_L2 | Tukey's post hoc<br>D4_L2 | Tukey's post hoc<br>D3_L2 v D4_L2 |
| Group effect | F (2, 31) = 0.8442 | P=0.4396 | Platform v Stimulation | Platform v Stimulation | Platform |
|  |  |  | q=2.492 ; Padj=0.1910 | q=2.902 ; Padj=0.1085 | q=3.919 ; Padj=0.0094 |
| Day effect | F (1, 31) = 19.31 | P=0.0001 | Platform v Food | Platform v Food | Stimulation |
|  |  |  | q= 4.995 ; Padj=0.0022 | q= 3.746 ; Padj=0.0272 | q= 1.995 ; Padj=0.1683 |
| Interaction | F (2, 31) = 6.224 | P=0.0053 | Stimulation v Food | Stimulation v Food | Food |
|  |  |  | q= 1.378 ; Padj=0.5958 | q= 0.000 ; Padj>0.9999 | q= 5.971 ; Padj=0.0002 |
| Speed<br>2-way RM ANOVA | F(DFn, DFd) | P-value | Tukey's post hoc<br>D3_L2 | Tukey's post hoc<br>D4_L2 | Tukey's post hoc<br>D3_L2 v D4_L2 |
| Group effect | F (2, 31) = 13.18 | P<0.0001 | Platform v Stimulation | Platform v Stimulation | Platform |
|  |  |  | q=7.288 ; Padj<0.0001 | q=5.944 ; Padj=0.0003 | q=0.5002 ; Padj=0.7260 |
| Day effect | F (1, 31) = 18.22 | P=0.0002 | Platform v Food | Platform v Food | Stimulation |
|  |  |  | q= 3.502 ; Padj=0.0418 | q= 0.6392 ; Padj=0.8937 | q= 2.466 ; Padj=0.0911 |
| Interaction | F (2, 31) = 7.640 | P=0.0020 | Stimulation v Food | Stimulation v Food | Food |
|  |  |  | q= 4.575 ; Padj=0.0055 | q= 6.439 ; Padj<0.0001 | q= 8.300 ; Padj<0.0001 |
| Latency to exit the first alley<br>2-way RM ANOVA | F(DFn, DFd) | P-value | Tukey's post hoc<br>D3_L2 | Tukey's post hoc<br>D4_L2 | Tukey's post hoc<br>D3_L2 v D4_L2 |
| Group effect | F (2, 31) = 39.31 | P<0.0001 | Platform v Stimulation | Platform v Stimulation | Platform |
|  |  |  | q=12.53 ; Padj<0.0001 | q=9.363 ; Padj<0.0001 | q=0.1343 ; Padj=0.9250 |
| Day effect | F (1, 31) = 6.810 | P=0.0138 | Platform v Food | Platform v Food | Stimulation |
|  |  |  | q=1.685 ; Padj=0.4629 | q=1.000 ; Padj=0.7602 | q=4.447 ; Padj=0.0037 |
| Interaction | F (2, 31) = 3.724 | P=0.0355 | Stimulation v Food | Stimulation v Food | Food |
|  |  |  | q=11.23 ; Padj<0.0001 | q=8.589 ; Padj<0.0001 | q=1.024 ; Padj=0.4745 |
| Time spent in the new alley<br>2-way RM ANOVA | F(DFn, DFd) | P-value | Tukey's post hoc<br>D3_L2 | Tukey's post hoc<br>D4_L2 | Tukey's post hoc<br>D3_L2 v D4_L2 |
| Group effect | F (2, 31) = 8.802 | P=0.0009 | Platform v Stimulation | Platform v Stimulation | Platform |
|  |  |  | q=3.919 ; Padj=0.0199 | q=2.373 ; Padj=0.2216 | q=1.983 ; Padj=0.1709 |
| Day effect | F (1, 31) = 29.92 | P<0.0001 | Platform v Food | Platform v Food | Stimulation |
|  |  |  | q=7.337 ; Padj<0.0001 | q=3.043 ; Padj=0.0879 | q=3.407 ; Padj=0.0221 |
| Interaction | F (2, 31) = 6.015 | P=0.0062 | Stimulation v Food | Stimulation v Food | Food |
|  |  |  | q=1.764 ; Padj=0.4301 | q=0.01628 ; Padj>0.9999 | q=8.916 ; Padj<0.0001 |

| Level 3 |  |  |  |  |
| --- | --- | --- | --- | --- |
| Distance<br>2-way RM ANOVA | F(DFn, DFd) | P-value | D4_L3<br>Tukey's post hoc |  |
|  |  |  | S3 | S4 |
| Group effect | F (2, 31) = 13.13 | P<0.0001 | Platform v Stimulation<br>q=3.310 ; Padj=0.1123 | Platform v Stimulation<br>q=1.071 ; Padj=0.7383 |
| Session effect | F (6.342, 196.6) = 11.52 | P<0.0001 | Platform v Food<br>q= 0.4571 ; Padj=0.9442 | Platform v Food<br>q= 3.291 ; Padj=0.0793 |
| Interaction | F (26, 403) = 2.842 | P<0.0001 | Stimulation v Food<br>q= 3.111 ; Padj=0.1411 | Stimulation v Food<br>q= 2.975 ; Padj=0.1772 |
| D5_L3<br>Tukey's post hoc | S1 | S2 | S3 | S4 |
|  | Platform v Stimulation<br>q=4.728 ; Padj=0.0358 | Platform v Stimulation<br>q=3.281 ; Padj=0.1302 | Platform v Stimulation<br>q=4.373 ; Padj=0.0488 | Platform v Stimulation<br>q=6.011 ; Padj=0.0110 |
|  | Platform v Food<br>q= 4.355 ; Padj=0.0168 | Platform v Food<br>q= 2.720 ; Padj=0.1522 | Platform v Food<br>q= 1.277 ; Padj=0.6434 | Platform v Food<br>q= 1.935 ; Padj=0.3725 |
|  | Stimulation v Food<br>q= 1.392 ; Padj=0.6027 | Stimulation v Food<br>q= 2.221 ; Padj=0.3289 | Stimulation v Food<br>q= 3.635 ; Padj=0.0861 | Stimulation v Food<br>q= 4.777 ; Padj=0.0267 |
| D6_L3<br>Tukey's post hoc | S1 | S2 | S3 | S4 |
|  | Platform v Stimulation<br>q=2.522 ; Padj=0.2645 | Platform v Stimulation<br>q=6.042 ; Padj=0.0076 | Platform v Stimulation<br>q=4.505 ; Padj=0.0432 | Platform v Stimulation<br>q=1.000 ; Padj=0.7623 |
|  | Platform v Food<br>q= 5.080 ; Padj=0.0057 | Platform v Food<br>q= 0.2210 ; Padj=0.9866 | Platform v Food<br>q= 2.062 ; Padj=0.3279 | Platform v Food<br>q= 0.08698 ; Padj=0.9979 |
|  | Stimulation v Food<br>q= 1.138 ; Padj=0.7150 | Stimulation v Food<br>q= 5.752 ; Padj=0.0084 | Stimulation v Food<br>q= 3.421 ; Padj=0.1064 | Stimulation v Food<br>q= 1.309 ; Padj=0.6369 |
| D7_L3<br>Tukey's post hoc | S1 | S2 | S3 | S4 |
|  | Platform v Stimulation<br>q=4.555 ; Padj=0.0476 | Platform v Stimulation<br>q=4.367 ; Padj=0.0376 | Platform v Stimulation<br>q=5.972 ; Padj=0.0150 | Platform v Stimulation<br>q=4.367 ; Padj=0.0394 |
|  | Platform v Food<br>q= 4.761 ; Padj=0.0116 | Platform v Food<br>q= 1.225 ; Padj=0.6671 | Platform v Food<br>q= 3.621 ; Padj=0.0562 | Platform v Food<br>q= 3.284 ; Padj=0.0778 |
|  | Stimulation v Food<br>q=0.5537 ; Padj=0.9195 | Stimulation v Food<br>q= 4.006 ; Padj=0.0684 | Stimulation v Food<br>q=0.4025 ; Padj=0.9564 | Stimulation v Food<br>q=0.8100 ; Padj=0.8365 |

| Level 3 |  |  |  |  |  |
| --- | --- | --- | --- | --- | --- |
| Distance change overnight<br>2-way *RM ANOVA* | F(DFn, DFd) | P-value | Tukey's post hoc<br>D4_L3-D5_L3 | Tukey's post hoc<br>D5_L3-D6_L3 | Tukey's post hoc<br>D6_L3-D7_L3 |
| Group effect | F (2, 31) = 16.86 | P<0.0001 | Platform v Stimulation<br>q=5.644 ; Padj=0.0028 | Platform v Stimulation<br>q=0.01393 ; Padj>0.9999 | Platform v Stimulation<br>q=4.161 ; Padj=0.0259 |
| Day effect | F (1.579, 48.96) = 0.09157 | P=0.8695 | Platform v Food<br>q= 6.544 ; Padj=0.0003 | Platform v Food<br>q= 3.227 ; Padj=0.0859 | Platform v Food<br>q= 3.713 ; Padj=0.0377 |
| Interaction | F (4, 62) = 3.314 | P=0.0159 | Stimulation v Food<br>q= 2.247 ; Padj=0.2758 | Stimulation v Food<br>q= 1.305 ; Padj=0.6450 | Stimulation v Food<br>q=0.1810 ; Padj=0.9910 |
| One sample Wilcoxon test | D4_L3-D5_L3 | D5_L3-D6_L3 | D6_L3-D7_L3 |  |  |
| Platform | W=-83 ; P=0.0067 | W=-5 ; P=0.9032 | W=-85 ; P=0.0052 |  |  |
| Stimulation | W=15 ; P=0.1562 | W=-9 ; P=0.4375 | W=19 ; P=0.0625 |  |  |
| Food | W=77 ; P=0.0134 | W=73 ; P=0.0203 | W=57 ; P=0. 0.0785 |  |  |
| Within-day distance change<br>2-way *RM ANOVA* | F(DFn, DFd) | P-value | Tukey's post hoc<br>D5_L1 | Tukey's post hoc<br>D6_L3 | Tukey's post hoc<br>D7_L3 |
| Group effect | F (2, 31) = 2.614 | P=0.0893 | Platform v Stimulation<br>q=1.684 ; Padj=0.4963 | Platform v Stimulation<br>q=2.502 ; Padj=0.2500 | Platform v Stimulation<br>q=2.783 ; Padj=0.1811 |
| Day effect | F (1.964, 60.90) = 9.850 | P=0.0002 | Platform v Food<br>q=3.916 ; Padj=0.0309 | Platform v Food<br>q=3.178 ; Padj=0.0824 | Platform v Food<br>q=2.173 ; Padj=0.2988 |
| Interaction | F (4, 62) = 3.623 | P=0.0102 | Stimulation v Food<br>q=1.298 ; Padj=0.6402 | Stimulation v Food<br>q=0.9569 ; Padj=0.7848 | Stimulation v Food<br>q=0.04930; Padj=0.9993 |
| One sample Wilcoxon test | D5_L3 | D6_L3 | D7_L3 |  |  |
| Platform | W=-49 ; P=0.1353 | W=23 ; P=0.5016 | W=43 ; P=0.1885 |  |  |
| Stimulation | W=-13 ; P=0.2188 | W=-17 ; P=0.0938 | W=21 ; P=0.0312 |  |  |
| Food | W=-95 ; P=0.0134 | W=-71 ; P=0.0203 | W=-57 ; P=0. 0.0785 |  |  |

**Supplementary Table 1: Statistical analyses for all levels performances across the three reward conditions (MFB stimulation, food, and platform escape).**
| Level 3 |  |  |  |  |  |  |  |
| --- | --- | --- | --- | --- | --- | --- | --- |
| Percentage of direct trials<br>2-way *RM ANOVA* | F(DFn, DFd) | P-value | Tukey's post hoc<br>D4_L3 | Tukey's post hoc<br>D5_L3 | Tukey's post hoc<br>D6_L3 | Tukey's post hoc<br>D7_L3 | Tukey's post hoc<br>D4_L3 v D7_L3 |
| Group effect | F (2, 31) = 1.437 | P=0.2531 | Platform v Stimulation<br>q=0.9070 ; Padj=0.8025 | Platform v Stimulation<br>q=1.557 ; Padj=0.5312 | Platform v Stimulation<br>q=1.286 ; Padj=0.6446 | Platform v Stimulation<br>q=1.651 ; Padj=0.4885 | Platform<br>q=8.054 ; Padj=0.0004 |
| Day effect | F (2.724, 84.46) = 29.11 | P<0.0001 | Platform v Food<br>q= 2.346 ; Padj=0.2409 | Platform v Food<br>q= 1.207 ; Padj=0.6742 | Platform v Food<br>q= 0.6242 ; Padj=0.8987 | Platform v Food<br>q= 4.097 ; Padj=0.0206 | Stimulation<br>q= 4.375 ; Padj=0.0920 |
| Interaction | F (6, 93) = 1.124 | P=0.3544 | Stimulation v Food<br>q= 0.4345 ; Padj=0.9497 | Stimulation v Food<br>q= 0.6629 ; Padj=0.8874 | Stimulation v Food<br>q= 0.9153 ; Padj=0.7988 | Stimulation v Food<br>q= 2.898 ; Padj=0.1293 | Food<br>q= 8.060 ; Padj=0.0004 |
| Speed<br>2-way *RM ANOVA* | F(DFn, DFd) | P-value | Tukey's post hoc<br>D4_L3 | Tukey's post hoc<br>D5_L3 | Tukey's post hoc<br>D6_L3 | Tukey's post hoc<br>D7_L3 | Tukey's post hoc<br>D4_L3 v D7_L3 |
| Group effect | F (2, 31) = 24.29 | P<0.0001 | Platform v Stimulation<br>q=6.359 ; Padj=0.0099 | Platform v Stimulation<br>q=6.501 ; Padj=0.0075 | Platform v Stimulation<br>q=3.696 ; Padj=0.0817 | Platform v Stimulation<br>q=1.490 ; Padj=0.5754 | Platform<br>q=1.013 ; Padj=0.8889 |
| Day effect | F (2.793, 86.60) = 42.64 | P<0.0001 | Platform v Food<br>q= 0.4940 ; Padj=0.9352 | Platform v Food<br>q= 5.901 ; Padj=0.0019 | Platform v Food<br>q= 7.543 ; Padj=0.0001 | Platform v Food<br>q= 9.528 ; Padj<0.0001 | Stimulation<br>q= 7.785 ; Padj=0.0101 |
| Interaction | F (6, 93) = 20.56 | P<0.0001 | Stimulation v Food<br>q= 5.911 ; Padj=0.0057 | Stimulation v Food<br>q= 9.330 ; Padj<0.0001 | Stimulation v Food<br>q= 8.394 ; Padj=0.0002 | Stimulation v Food<br>q= 6.276 ; Padj=0.0055 | Food<br>q= 15.46 ; Padj<0.0001 |
| Latency to exit the first alley<br>2-way *RM ANOVA* | F(DFn, DFd) | P-value | Tukey's post hoc<br>D4_L3 | Tukey's post hoc<br>D5_L3 | Tukey's post hoc<br>D6_L3 | Tukey's post hoc<br>D7_L3 | Tukey's post hoc<br>D4_L3 v D7_L3 |
| Group effect | F (2, 31) = 25.46 | P<0.0001 | Platform v Stimulation<br>q=3.937 ; Padj=0.0838 | Platform v Stimulation<br>q=4.504 ; Padj=0.0538 | Platform v Stimulation<br>q=5.438 ; Padj=0.0271 | Platform v Stimulation<br>q=4.401 ; Padj=0.0581 | Platform<br>q=0.8048 ; Padj=0.9395 |
| Day effect | F (1.232, 38.18) = 14.71 | P=0.0002 | Platform v Food<br>q=2.486 ; Padj=0.2088 | Platform v Food<br>q=5.072 ; Padj=0.0068 | Platform v Food<br>q=3.379 ; Padj=0.0614 | Platform v Food<br>q=0.4997 ; Padj=0.9337 | Stimulation<br>q=3.283 ; Padj=0.2121 |
| Interaction | F (6, 93) = 10.15 | P<0.0001 | Stimulation v Food<br>q=3.865 ; Padj=0.0887 | Stimulation v Food<br>q=4.191 ; Padj=0.0681 | Stimulation v Food<br>q=5.274 ; Padj=0.0304 | Stimulation v Food<br>q=4.372 ; Padj=0.0594 | Food<br>q=2.749 ; Padj=0.2580 |
| Time spent in the new alley<br>2-way *RM ANOVA* | F(DFn, DFd) | P-value | Tukey's post hoc<br>D4_L3 | Tukey's post hoc<br>D5_L3 | Tukey's post hoc<br>D6_L3 | Tukey's post hoc<br>D7_L3 | Tukey's post hoc<br>D4_L3 v D7_L3 |
| Group effect | F (2, 31) = 8.924 | P=0.0009 | Platform v Stimulation<br>q=2.946 ; Padj=0.1848 | Platform v Stimulation<br>q=4.531 ; Padj=0.0499 | Platform v Stimulation<br>q=2.343 ; Padj=0.3006 | Platform v Stimulation<br>q=3.077 ; Padj=0.1515 | Platform<br>q=6.854 ; Padj=0.0016 |
| Day effect | F (2.440, 75.65) = 16.58 | P<0.0001 | Platform v Food<br>q=2.378 ; Padj=0.2355 | Platform v Food<br>q=2.755 ; Padj=0.1587 | Platform v Food<br>q=1.963 ; Padj=0.3617 | Platform v Food<br>q=3.458 ; Padj=0.0637 | Stimulation<br>q=3.420 ; Padj=0.1906 |
| Interaction | F (6, 93) = 3.709 | P=0.0024 | Stimulation v Food<br>q=2.230 ; Padj=0.3289 | Stimulation v Food<br>q=2.851 ; Padj=0.1742 | Stimulation v Food<br>q=1.753 ; Padj=0.4793 | Stimulation v Food<br>q=0.1312 ; Padj=0.9953 | Food<br>q=5.558 ; Padj=0.0082 |

## Methods

### Experimental animals

Thirty-four male C57BL/6 J mice (8–14 weeks old, Janvier France) were used in this study. They were all trained in the Starmaze navigation task but distributed in three groups based on the modality of reward: in group 1 (n = 14), mice were trained in an aquatic version of the starmaze in which rewards consisted of a safe escape on a dry platform; in group 2 (n = 14), mice were food-deprived and rewards consisted of food pellets delivered through a food dispenser; in group 3 (n = 6) rewards were delivered through medial forebrain bundle (MFB) stimulation. Mice were housed in groups of up to 5 animals under standard conditions: 12 h light/dark cycle, with free access to water and food. Seven days prior to the beginning of behavioral tests, mice were isolated and housed in an animal facility in individual cages with access to enrichment (nesting material and a toy), and food deprivation was performed in group 2. All behavioral experiments took place during the light cycle and were conducted in accordance with the European Communities Council Directive (86/809/EEC) regarding the care and use of animals for experimental procedures in compliance with the *Ministère de l’Agriculture et de la Forêt, Service Vétérinaire de la Santé et de la Protection Animale* (#28025-2020111721361293 v8) of the Ministère de l’enseignement supérieur, de la recherche et de l’innovation and the approval of the ethical committees of Sorbonne Université (CEEA n°5).

To assess potential motor deficits following implantation, all three groups’ motor abilities were tested using a static balance and a motor coordination task.

#### Unstable platform

The static balance of mice was assessed using a 9 cm circular platform fixed on a 105 cm vertical rod. The platform tilted in all directions according to the position of the mouse. The holding time is recorded during three trials of 3 minutes maximum. If the mice fell before 20 seconds, they were immediately returned to the platform.

#### Accelerated Rotarod

Motor coordination was assessed using a rotating horizontal cylinder (44cm height). Mouse holding time was recorded during 3 trials. Each trial lasted a maximum of 5 minutes during which the speed was gradually increased from 4 to 40 rpm, with an increment of 1 rpm every 8 seconds.

### MFB implantation

Group 3 mice were implanted under deep isoflurane anesthesia with bipolar stimulation electrodes (two 150-µm-diameter stainless steel electrodes twisted together; PlasticsOne) targeting the left MFB (AP +1.4, ML +1.2, DV +4.8). After surgery, mice were allowed a minimum of 1 week of recovery before behavioral assays.

#### MFB functional implantation testing

To check for functional implantation of the MFB stimulating electrode, group 3 mice were tested in a place preference task using a non-transparent 31x24x40 cm box with two 2 cm diameter holes on the same wall (supp. Figure 1). Exploration of the left hole was automatically detected by Smart tracking software and was rewarded with an MFB electrical stimulation, while there was no consequence to exploring the right hole. Each stimulation consisted of a 140 Hz train of negative pulses lasting 100 ms. Stimulation intensity was incrementally increased from 0 to 40 µA, and two sessions of 180s for each intensity were performed over two days. MFB implantation was considered successful when mice reached a number of entries in the rewarded zone greater than twice the mean number of entries at 0 µA. The optimal intensity ranged from 30 to 40 uA, depending on individual mice.

#### Food deprivation

Before starting the training procedure in the Starmaze, group 2 mice were food-restricted for seven consecutive days. During this period, 2 to 4g of food was delivered daily to each mouse in order to maintain the weight loss below 15% of the mouse’s initial weight.

#### Sequential navigation in the Starmaze

In all mice, a sequence-based, reward-oriented navigation task was performed in the Starmaze composed of a central stem from which two Y-mazes originate (Figure 1). External and internal alleys were 40 cm and 24 cm long, respectively, and both were 18 cm wide. Low light intensity was maintained around 10-20 lux. White noise (50 dB) was broadcast in the testing facility to cover auditory cues that the mice could have used to orient themselves. A circular black curtain was placed around the maze so that the mice could not rely on visual cues to learn and memorize the path. For the aquatic starmaze, alleys were filled with water made opaque with an inert nontoxic product (Accuscan OP 301, Brenntage, Lyon, France) to hide a platform 1 cm below the surface in the aquatic version. Water temperature was maintained between 18 and 20°C.

On each trial, mice had to reach a goal zone from a departure point to receive a reward consisting either of MFB stimulation (3 trains of 100ms at 140Hz, 2 seconds apart) (starmaze with MFB), 3 food pellets (food-reward Starmaze), or water escape by setting on a dry platform (aquatic starmaze). The current intensity of the MFB stimulation was set between 20 and 30 µA at the beginning, but could be adjusted progressively to up to 50 µA depending on the motivation of the mouse.

The departure point and the goal zone were fixed so that the shortest way to reach the reward was to execute a left-right-left sequence of turns. Each trial lasted at most 90 seconds, with a 1-minute inter-trial period, during which the maze was cleaned using 20% ethanol in the dry Starmaze versions. The animal repeated four such trials within a session, and four sessions were performed per day, with a 30-minute inter-session period. This schedule was carried out for up to seven days.

As all three intersections of the maze were visually identical, mice had to remember the temporal sequence of turns to solve the task.

During the first three and a half days, mice were “pre-trained” in a progressively more complex maze configuration. During the first day, part of the alleys between the departure point and the goal location were closed, so the mice could only use one path to get to the reward. The following days (2, 3, and 4), alleys were progressively opened one after the other (Figure 2). To consolidate the learning of the task, the starmaze configuration was the same during the first two sessions of each day as during the last two sessions of the previous day.

For each session, the percentage of direct trials, latency to reward, latency to exit the departure alley, and the mean speed were measured.

##### Tissue processing and histological procedures

Following the final round of behavioral tests, animals in the MFB group were deeply anesthetized with an overdose of anesthetics (xylazine 20 mg/kg/ketamine 200 mg/kg, administered intraperitoneally) and perfused intracardially with saline, followed by 4% paraformaldehyde (PFA) in a 0.1M phosphate buffer (PB). The brains were removed, post-fixed for 2–3 days in 4% PFA, and then stored at 4°C in 0.1 M PB with 0.02% sodium azide.

For histological verification of the placement of the MFB electrode, the brains were embedded in 3% agarose and then sliced into 100 μm-thick coronal sections. These sections were mounted on slides and coverslipped with a mounting medium containing DAPI (Fluoromount-G™ with DAPI, Fisher Scientific). Images were taken using an Axiozoom v16 microscope (Carl Zeiss, France) equipped with a 2.3X objective. The tip of the electrode tracks was identified by localized mechanical lesions in the MFB according to a stereotaxic atlas (Franklin and Paxinos, 2007).

### Data acquisition and statistical analysis

Data acquisition was performed by means of a video recording system and the tracking software Anymaze.

Statistical analyses were run using GraphPad Prism version 9. Normality was tested using the Shapiro-Wilk test. An ordinary one-way ANOVA was performed for accelerated rotarod analysis. The Kruskal-Wallis test was performed for static balance analysis. Two-way repeated measures ANOVAs were performed for Starmaze performance analysis. The significance level was set at 5%.

## Acknowledgments.

We gratefully acknowledge the animal facility staff from Institut de Biologie Paris Seine for their support. Funding: This work was supported by ANR-18-CE16-0010-02 (RewInhib) (L.R.-R. and C.A.), (Pomaret-Delalande-FRM Phd fellowship (J.A.), Centre National de Recherche Scientifique (CNRS), Institut National de la Santé de et de la Recherche Médicale (Inserm) and Sorbonne Université. We thank Aurelie Watilliaux for helpful support in ethical procedures and finalization of the paper and all members of the Cerebellum, Navigation and Memory (CeZaMe) team for their insightful discussions.

## Author contributions

Research design: CR and LRR; Experimentation: CL and JA; Data Analysis: CA, CL and JA; Validation: CR, LRR; Writing First Draft, Revision, Review, and Editing: JA, CL, CA, CR, LRR; Funding, PhD supervision, Project management, reagents and resources: CA, CR and LRR.

